# The macroecology of immunity: predominant influence of climate on invertebrate immune response

**DOI:** 10.1101/2025.06.16.659911

**Authors:** Adam Z. Hasik, Maggie Blondeau, Jake Harvey, Tania Groleau, Tonia de Bellis, Eric J. Pedersen, Alex Córdoba-Aguilar, Katie Marshall, Laura Ferguson, Jean-Philippe Lessard

## Abstract

The immune system is the primary defense against parasites. With the ever-increasing rate of disease, epidemiologic models considering geographic variation in immune responses could prove useful. Despite increasing interest in the macroecology of parasitism and infectious diseases, we know little about the macroecology of immune responses. Host characteristics, parasite exposure, and environmental factors can all affect immunity, but how these factors interact to shape spatial variation in the strength of immune responses remains unexplored. We captured odonates (dragonflies and damselflies) and their conspicuous ectoparasitic mites across a geographic area spanning the temperate and boreal forest biomes in eastern Canada. We then conducted immune response bioassays on 1,237 individuals from 63 odonate species. We used linear regressions and structural equation models to relate immune responses to host body size, parasite load, pH, temperature, and precipitation while accounting for evolutionary relationships among host species. We found significant differences in the strength of immune response among host individuals, and this variation was best explained by climatic conditions, specifically decreasing with precipitation and, to a lesser degree, temperature. While host species significantly differed in immune response strength, we found no effect of host body size, evolutionary relationships among hosts, or parasitism on immune response. Our study investigating the drivers of immune response across dozens of species spread across two biomes is the most comprehensive to date. Climatic conditions have a strong influence on host immune response, regardless of host characteristics or parasitism rates. In this specific case, strong immune responses were associated with low levels of annual precipitation, which could relate to the role of cuticular melanin content in desiccation resistance, and the melanin-based encapsulation response being a byproduct of this adaptation. A spatially-explicit understanding of the biological processes affecting immunity could improve epidemiological models of disease risk that inform disease management globally.

## Introduction

Predicting parasite, pathogen, and disease spread is increasingly relevant and challenging in a highly-connected world (Tsiotas & Tselios 2022). Epidemiologic models are used to predict patterns of disease spread and implement control measures. In this respect, the immune system emerges prominently as the first line of defense against attack by parasites and pathogens. Yet, the factors driving variation in immunity among individuals and populations are poorly-studied and rarely factored into epidemiologic models (Becker *et al*. 2019). Characteristics of the host, exposure to parasites or pathogens, and the abiotic environment can interact in complex ways to affect immunity (Sweeny & Albery 2022), but their interactions are challenging to elucidate (Johnson *et al*. 2019).

As the immune system is the primary line of defense against infection by parasites, pathogens, and disease, it is assumed to be costly in terms of fitness and should therefore lead to tradeoffs with life-history traits (e.g., fecundity, fertility, Albery *et al*. 2021). Although a plethora of studies have provided key evidence of immune variation due to such tradeoffs, most studies emphasize the role of biotic factors such as predation (Duong & McCauley 2016) and resource availability (Hasik *et al*. 2024b) without considering that of abiotic factors (Lazzaro & Little 2008). A relationship between immune response and temperature is expected in both invertebrate ectotherms (e.g., Mastore *et al*. 2019) and vertebrate endotherms (Butler *et al*. 2013), due to the thermal sensitivity of the enzymes involved in immune responses (e.g., Catalán *et al*. 2012). When one scales this temperature-dependent immunity to explore the effect of climate (specifically, temperature and humidity); climate is expected to be a clear driver of geographic variation in immunity (Li *et al*. 2024).

Parasites are a leading cause of disease and death around the world and thus are drivers of life history evolution via their effects on host fitness (Hasik & Siepielski 2022a) that have the potential to affect host macroevolutionary dynamics (Hasik *et al*. 2025). The majority of organisms on earth are infected by at least one parasite (Price 1980), and yet, we have a very limited understanding of the multifarious factors governing the intensity of infection and therefore, the health cost. Both among-individual and interspecific variation in immune response surely plays a central role, but the factors regulating immunity in natural settings are poorly understood, which can interfere with the accuracy of predictive epidemiologic models. Environmental factors and local parasite pressure can independently drive differences in immunity across space, but they could also act in concert (Becker *et al*. 2020). Parasitism varies among host populations distributed across large-scale environmental gradients (LoScerbo *et al*. 2020; Hasik & Siepielski 2022b) and at fine spatial scales, within populations (Albery *et al*. 2019; Hasik *et al*. 2024b). To date, however, the focus on a limited set of taxa, specifically vertebrates (Becker *et al*. 2020), limits our ability to identify generalities regarding the relative influence of environmental conditions and parasitism on immune defenses that would apply across host-parasite systems (Rolff & Siva-Jothy 2003).

Here, we use data from a large-scale field sampling of communities of dragonflies and damselflies and their larval water mite ectoparasites to assess the relative influence of the abiotic environment, parasite pressure, and host characteristics on host immunity. We quantified the magnitude of melanin encapsulation, which is a well-studied aspect of invertebrate immunity, and related among-host individual variation in this immunological trait to variation in host body size, parasitism, and the abiotic environment (pH, temperature, precipitation). We then sought to disentangle the complex and interconnected network of relationships between these factors using structural equation models (SEMs). Specifically, we predicted that the local abiotic environment and parasitism (itself influenced by the local environment) would act together to drive variation in immune defenses. While our focus is understanding if and how the local abiotic environment drives immune defenses over a large geographic area, there is the possibility that immune function is an inherent property of the host species and varies independent of the biotic or abiotic environment. Likewise, other traits of interest (i.e., host size, mite parasitism) may also be independent properties of the host species. Therefore, we also measured the strength of the phylogenetic signal of immunity, parasitism, and host size.

## Methods

### Study system

For our hosts we collected odonates (e.g., dragonflies and damselflies); invertebrates that spend the majority of their lives in the water column as aquatic larvae. Adult odonates are frequently parasitized by larval *Arrenurus* water mites (Acari: Hydrachnida) which leave the host when the latter returns to water to reproduce as adults (Smith *et al*. 2010). Larval water mites are opportunistic parasites, and a single species can parasitize multiple odonate species (Zawal & Buczyński 2013), while a single odonate species can be simultaneously parasitized by several mite species (Smith & Oliver 1986). Water mite parasitism is affected by both host traits such as body weight and environmental factors such as pH, conductivity, dissolved oxygen, and air temperature (LoScerbo *et al*. 2020; Hasik *et al*. 2024a). There is also substantial variation in mite prevalence between and within species (Ilvonen *et al*. 2018; Hasik & Siepielski 2022b; Hasik *et al*. 2024a) and water mites reduce host survival (Forbes & Baker 1991), although there appears to be no general trend toward either sex being more infected (Ilvonen *et al*. 2016). Both climatic variables and water chemistry influence not only odonate community composition in aquatic systems, but also parasite prevalence and intensity (Arrowsmith *et al*. 2018; LoScerbo *et al*. 2020; Hasik & Siepielski 2022b; Hasik *et al*. 2024a).

Odonates defend themselves from water mites via a general immune defense shared among invertebrates: the phenoloxidase cascade (González-Santoyo & Córdoba-Aguilar 2012). In short, the phenoloxidase cascade produces melanin as its end product, which encases water mite feeding tubes and prevents the mites from gorging on the host’s haemolymph (Hasik *et al*. 2023). Though it is a general defensive mechanism, resistance to water mites is known to vary intra- and inter-specifically. Resistance is greater when the host is locally rare (Lajeunesse *et al*. 2004), increases with both the prevalence and intensity of parasitism (Nagel *et al*. 2010; Kaunisto *et al*. 2017), the degree of local adaptation to parasites (Gómez-Llano *et al*. 2020), and resource availability (Hasik *et al*. 2021).

### Odonate sampling

We opportunistically sampled odonates using a butterfly net in the summers of 2019 and 2020. We standardized the time of day for all sampling events and found no evidence for an effect of temperature at time of collection or date (i.e., seasonal effects) on immune responses (Fig. S1). We collected animals near water bodies at 42 sites ranging from the northern temperate forest region to the middle of the boreal forest in southeastern Canada (Fig. 1). See Table S1 for detailed information on sample sizes and species compositions. In total, we collected 1,237 individuals from 63 species broadly distributed across the odonate phylogenetic tree.

**Fig 1.**
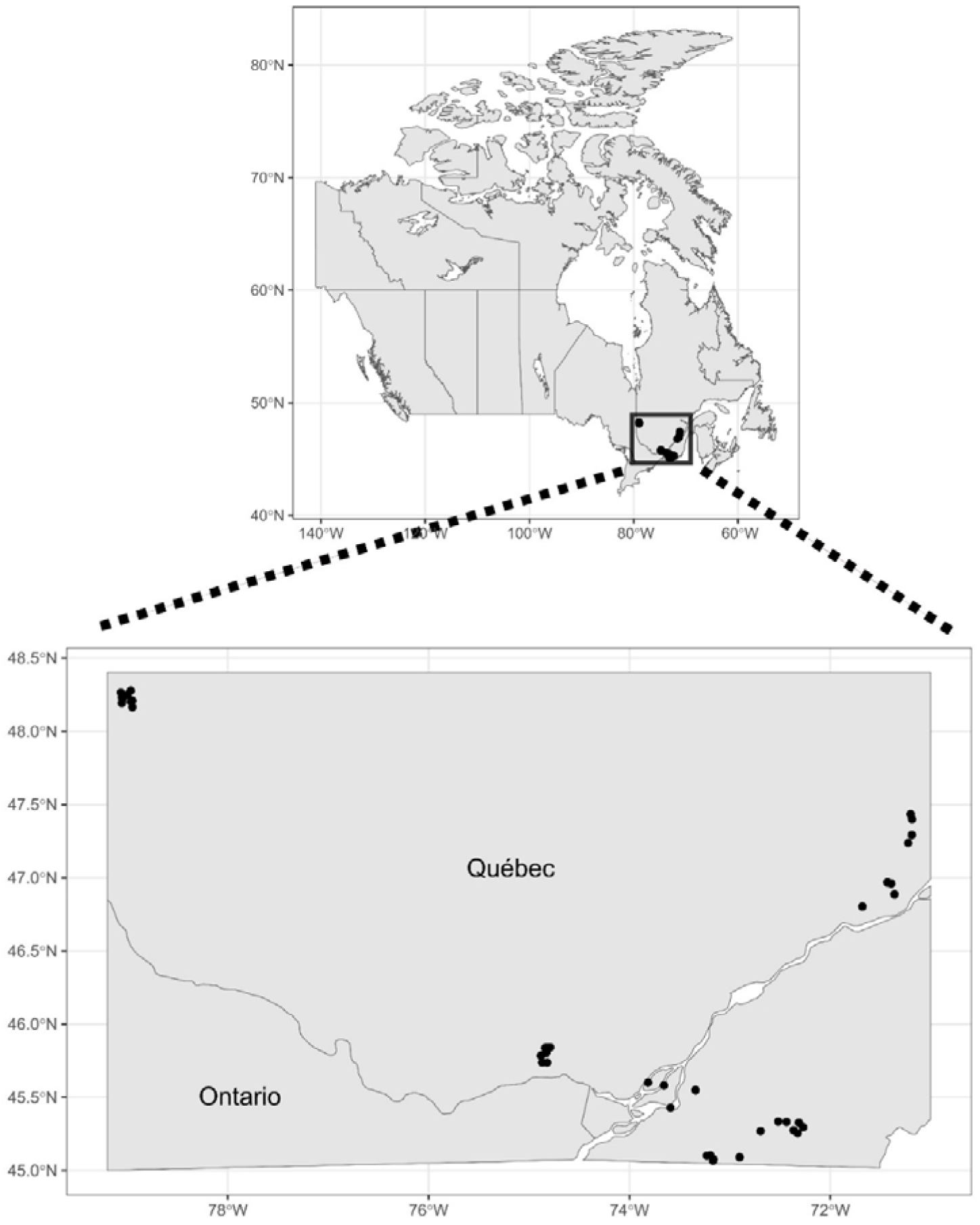
Map of the study sites across southeastern Canada, with the lower panel showing a zoomed in perspective of the sites spread across Québec.

### Immune response

Melanin, particularly eumelanin, is the encapsulating defense molecule and is dark brown in color (González-Santoyo & Córdoba-Aguilar 2012), so by quantifying the darkness of a proxy that has been inserted and melanized within a live odonate (to simulate a mite feeding tube), we can determine the relative strength of its immune response (Ilvonen & Suhonen 2016).

For full details of the immune assays see the Supplemental Materials. Briefly, we inserted a sterilized 3mm piece of nylon monofilament (hereafter implant) into the venter of the odonate thorax. This region of the body was chosen because it is where mites tend to latch onto their host, and the implant is meant to imitate the feeding tube of the parasite (Hasik *et al*. 2023). After 12 hours we removed and photographed the implant directly above (∼30cm between implant and camera lens, centering the lens over the implant) using the flash from the camera (Canon EOS Rebel T6s, lens extension MP-E 65mm). We then photographed the complete odonate animal, sacrificed them with acetone, and then stored them in glassine envelopes for subsequent identification and mite counting.

### Quantifying immune response

We used ImageJ (Schneider *et al*. 2012) to quantify the degree of melanization of each implant. We converted images of the nylon implant to 8-bit greyscale, which allowed us to measure average grey value of the pixels. Converting images to greyscale also allows us to extract darkness values ranging from 0 (completely black) to 255 (completely white), which we used to obtain the value representing immune response strength.

We selected six squares on each implant that had dimensions equal to the width of the nylon monofilament, ensuring that the areas were all proportional as every filament has the same width (0.4 mm) across images. The six squares that we measured were: (1) the deepest point of insertion, (2) the area nearest the point of insertion (just under the cuticle), (3) the area directly in between point 1 and 2, (4) the end of the side of the nylon not inserted into the specimen, (5) the region just outside of the cuticle that was not inserted, (6) and the region directly in the middle of 4 and 5 (Fig. S2). See the Supplemental Materials for further information on our use of six measurement points. We compared the average grey value of the three measurements spanning the inserted portion (the inserted average, denoted SA) to the average grey value of the three measurements spanning the portion that was not inserted were taken (the inserted control, denoted CA). The immune response for a given individual *i* was calculated by taking the difference between the control average for that individual (*CA_i_*) and the inserted average (*SA_i_*):

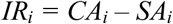

Note that higher values of *IR_i_* indicate a stronger melanization response, since *CA_i_* and *SA_i_* are grey-scale values where 0 represents a completely black value and 255 represents a white value. As such, the higher the melanization in the sample, the closer to zero the value for *SA_i_* will be.

### Host characteristics

Parasitism and immune function are known to vary with body size (Hasik & Siepielski 2022b). To accurately quantify body mass, we weighed each of the dried odonate host individuals using an electronic analytical scale. To mitigate specimen damage and prevent limb or fragment loss, specimens were removed from their glassine envelope with metal pliers and placed in small plastic square weighing boats. Furthermore, weighing boats were replaced between each measurement to minimize static interference, ensuring consistency in recorded mass.

### Parasitism

To quantify parasitism for each odonate host individual, we examined dry specimens under a dissecting microscope (LoScerbo *et al*. 2020). We counted the ectoparasitic mites that were attached to the host and noted the points of attachment on the host body in accordance with previous studies (Forbes *et al*. 1999). We included detached, engorged mites found within the individual odonate collection envelope in the total mite count for the host individual.

### Evolutionary relationships

We inferred evolutionary relationships among species using the time-calibrated odonate phylogeny of Waller and Svensson (2017), as it is the most comprehensive and well-resolved phylogeny for this group. Waller & Svensson’s tree was constructed using DNA sequences and contained 1,322 species (∼21% of all odonate species). We pruned this tree to include only those species which we collected during the field surveys using the *ape* package (Paradis *et al*. 2004) in R v4.1.2 (2021). Though our analyses included 63 total species, only 53 of those species have been mapped to the odonate phylogeny, thus we were not able to include all species in our phylogenetic analyses.

### Abiotic environment

We quantified the ambient climate and water pH at each collection site. We extracted mean annual temperature (°C; BIO1) and mean annual precipitation (mm; BIO12) associated with each lake from WorldClim version 2.1 (Hijmans *et al*. 2005) as climatic variables representing physiological constraints on hosts and parasites in our models. These data are calculated using climate data from 1970-2000. We recorded lake pH using an Oakton WD-35634-35 Series 50 Waterproof Pocket Ph/Cond/TDS/Salinity Tester. pH was quantified as the mean of five measurements were taken at least 5m apart where possible, and between 1 – 3m away from the shoreline.

### Statistical analyses

We conducted all statistical analyses in R v4.1.2 (2021). Our first question was concerned with understanding if the local environment and parasitism were associated with immune response over large spatial scales. To test this, we used our entire dataset (i.e., records from every species and population) to relate melanization response to environmental factors and parasitism. This dataset included observations of immune response for 1,237 individuals from 63 species. We used the **lme4** package (Bates 2010) to estimate a Gaussian linear mixed-effect model shown in Equation (1), using the mixed effect model notation suggested by Zuur and Leno (2016) where coefficients for fixed effect terms are implicit.

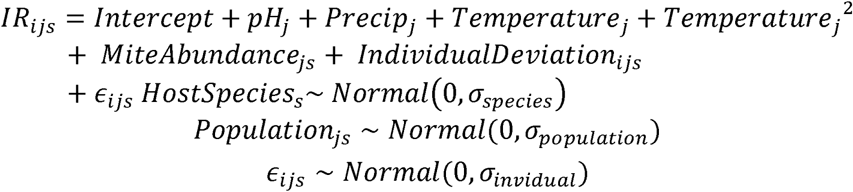

Here *IR_ijs_* is the immune response index for individual *i* of species *s* in site *j*, *pH_j_*, *Precip_j_*, and *Temperature_j_* are fixed effects of the three environmental variables (all variables were Z-score transformed in the full dataset before analysis). We also included *Temperature_j_*^2^ due to the non-linear relationship between temperature and melanization. *MiteAbundance_j_* and *IndividualDeviation_ijs_* refer to the population-level mean mite abundance for species *s* at site *j* and individual-level deviation from this mean for individual *i* of species *s* at site *j*, respectively. Using this modelling approach allows us to avoid confounding the effect of mite density *on the individual* with effects of *average mite abundance on individuals in the population.* An individual might have a higher immune response because it is living in a population that has a higher-than typical average level of mites (i.e., population effect of parasite load on *IR_ijs_*) or it might have a higher immune response because it has higher than average levels of parasites itself (or both effects could be present). It could also be that these effects could occur in opposite directions; adaptive evolution could result in *IR_ijs_* being stronger on average in areas with high mite abundances, but an individual with a large number of mites might have a reduced immune response if mite infection depleted their energy stores before our melanization test. *HostSpecies_s_* and *Population_js_* are species and population (species nested in site) specific random effects, and *E_ijs_* is residual individual-specific error. Random effects were assumed to have a normal distribution.

Our second question was concerned with disentangling the complex and interconnected network of relationships between the local environment, parasitism, and immunity. To do so we utilized structural equation models (SEMs, Shipley 2009). Fig. S3 contains a graphical representation of the initial SEM we used to test our hypothesis of water mite-mediated indirect relationships between the local aquatic environment and immune defenses. For this analysis we used a subset of our data, limiting it to those species for which we had roughly equivalent representation of infected and uninfected individuals with a minimum of *n* = 5 individuals from *n* = 5 populations, including observations of immune response for 556 individuals from seven species. We did this because we were interested in understanding potential drivers of parasitism and immunity within populations, thus it was important to include infected and uninfected individuals and remove those species that were rarely infected and had limited sampling. For all models we compared goodness-of-fit using the comparative fit index (CFI, a measure of the amount of variance accounted for in the covariance matrix, Fan et al., 2016) and root mean square error of approximation (a measure of how far a hypothesized model is from a model with a better fit, Xia & Yang 2018). Models with CFI > 0.95, and RMSE < 0.06 are considered to fit the data well and were compared to one another using AIC values (as in Hasik & Siepielski 2022b; Hasik *et al*. 2024a).

Our initial SEM tested the hypothesis that the local abiotic environment (pH) was affected by the local climate (precipitation and temperature), that host size was related to the local climate (precipitation and temperature), that parasitism population-level mean mite abundance was affected by the local environment (pH) and climate (precipitation), that individual-level deviance in infection was affected by the local abiotic environment (pH) and climate (precipitation) and host size, and finally that melanization response was affected by parasitism (population-level and individual-level), the local abiotic environment (pH), and climate (precipitation and temperature). There is, however, a possibility that there are connections between these various predictors that we did not model (i.e., temperature affects parasitism). To test for the presence of other relationships, we identified additional terms to fit to the model by analyzing modification indices using the “modindices” argument in **lavaan** (Rosseel 2012). Modification indices with χ*^2^* > 3.84 identify what terms, if added to a candidate model, improve model fit (Whittaker 2012). This method allowed us to build a statistically-informed network of additional relationships between the local abiotic environment, climate, parasitism, and melanization response. We iteratively added direct predictors of host size and mite abundance (population- and individual-level) with χ*^2^* > 3.84 until no further modification indices were identified and the model had a CFI > 0.95 and RMSE < 0.06 (as in Hasik *et al*. 2024a). We centered all environmental variables to have a mean of 0 and SD of 1 and included host species as a random effect.

### Phylogenetic signal

To assess the influence of evolutionary history on host traits, we conducted a test of phylogenetic signal. Specifically, we used Blomberg’s *K* to test for phylogenetic signal on host body weight, individual-level parasitism, and immune response (Blomberg *et al*. 2003). In short, Blomberg’s *K* estimates the strength of a phylogenetic signal compared to the expectations under Brownian motion, with *K* = 1 representing the expected value under Brownian motion and *K* < 1 and *K* > 1 representing a weaker or stronger signal than expected, respectively (Blomberg *et al*. 2003). We used the “phylosig” function from the **phytools** package (Revell 2012) to calculate *K* for species’ mean body weight, mean mite abundance, and mean melanization response. Here we estimated species mean mite abundance as the mean abundance of mites for all individuals of a given species, pooling individuals from all populations.

## Results

### The local climate predicts immune defenses

When considering our entire dataset, we found that melanization response strongly declined as mean annual precipitation of each site increased (χ*^2^* = 180.67, *p* < 0.0001) and weakly declined as mean annual temperature of each site increased (χ*^2^_1_* = 17.28, *p* < 0.0001, Fig. 2), though the quadratic term for temperature was not significant (χ*^2^* = 0.61, *p* = 0.43). We did not find significant relationships between melanization response and site pH (χ*^2^_1_* = 1.63, *p* = 0.20), population-level mite abundance (χ*^2^_1_* = 0.01, *p* = 0.94), or individual-level deviance in mite abundance (χ*^2^* = 1.01, *p* = 0.31).

**Fig 2.**
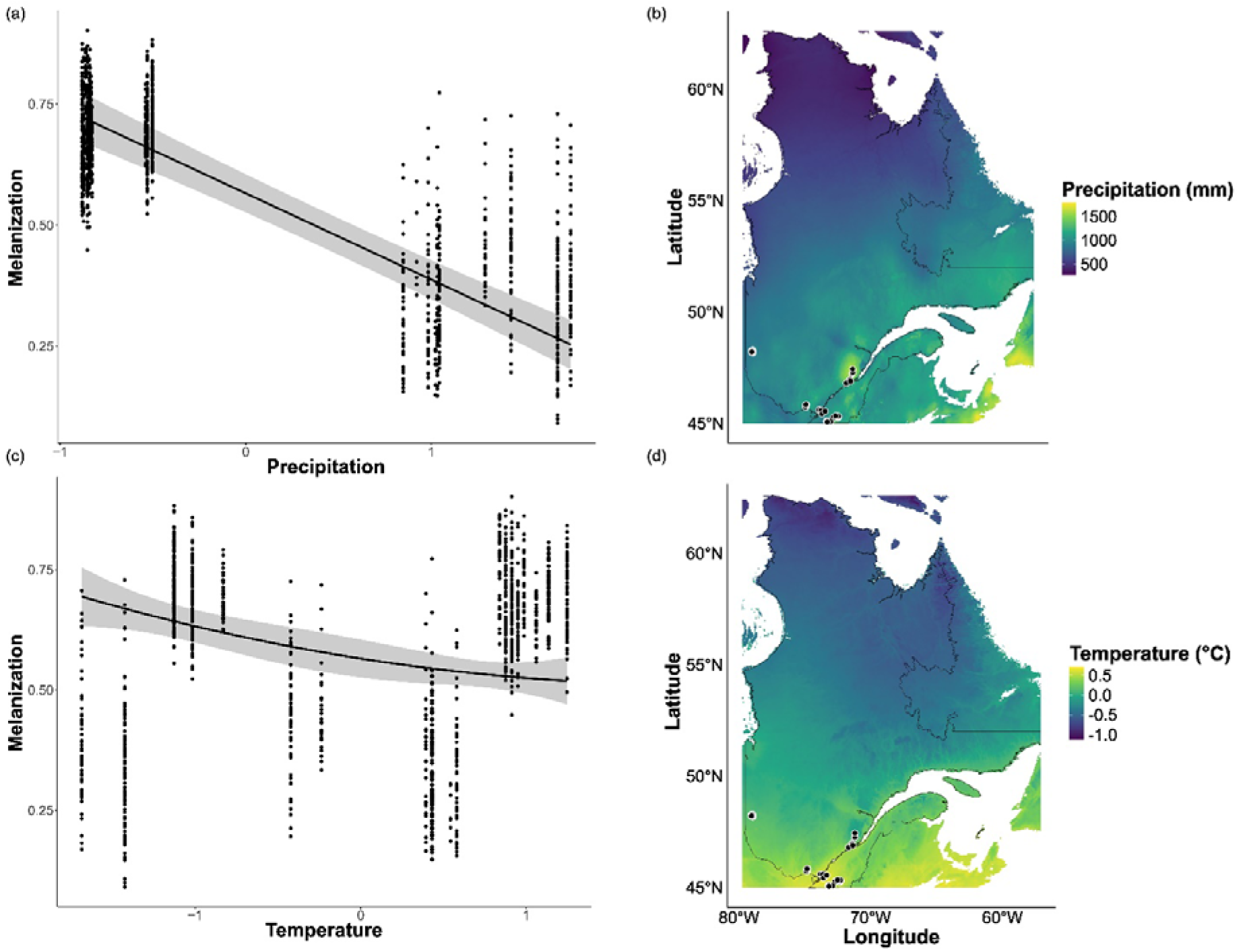
Relationships between melanization and environmental factors. Shown are model-derived predicted regressions (bands denote 95% CI) of melanization response on precipitation (a) and temperature (c), as well as climate maps of Québec denoting mean annual precipitation (b) and temperature (d) with points denoting sampling sites. Note the different x-axes between panels (a) and (c). Points denote melanization values for individual odonates.

### Environmental drivers of immunity

Our SEMs revealed that there were no indirect relationships among climate, pH, mite parasitism, and immune response. Instead, we found that melanization weakly increased with body weight and population mean mite abundance while declining moderately, strongly, and weakly with temperature, precipitation, and individual deviance, respectively (Fig. 3). Additionally, we found that pH moderately declined with precipitation (Fig. 3).

**Fig 3.**
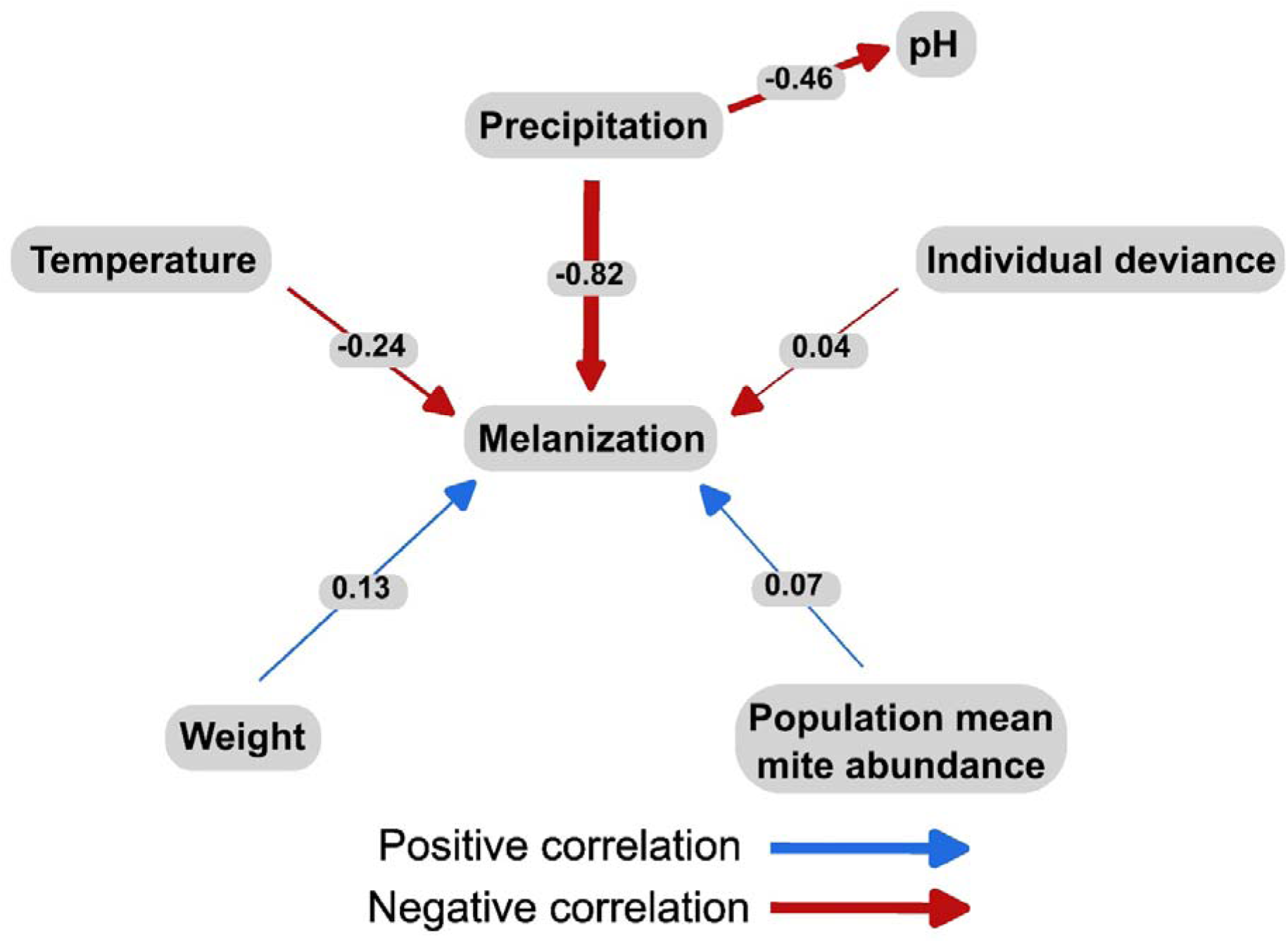
SEM plot of the relationships between climatic variables, the local environment, parasitism, and melanization. Only significant relationships are shown, with numbers representing standardized effect sizes. Width of the arrows denotes effect size magnitude, and color denotes direction of the relationship.

### Phylogenetic signal among traits

We found no evidence for a phylogenetic signal of mite abundance (*K* = 0.10, *p* = 0.47) or melanization (*K* = 0.04, *p* = 0.40), suggesting that neither one of these traits is phylogenetically constrained. However, we did find a strong and significant signal for host size (*K* = 0.99, *p* = 0.001, Fig. 4), suggesting that host body size is as expected under Brownian motion. That we did not find a signal for melanization is consistent with how wildly immune responses varied among host species and within genera (Fig. 5).

**Fig 4.**
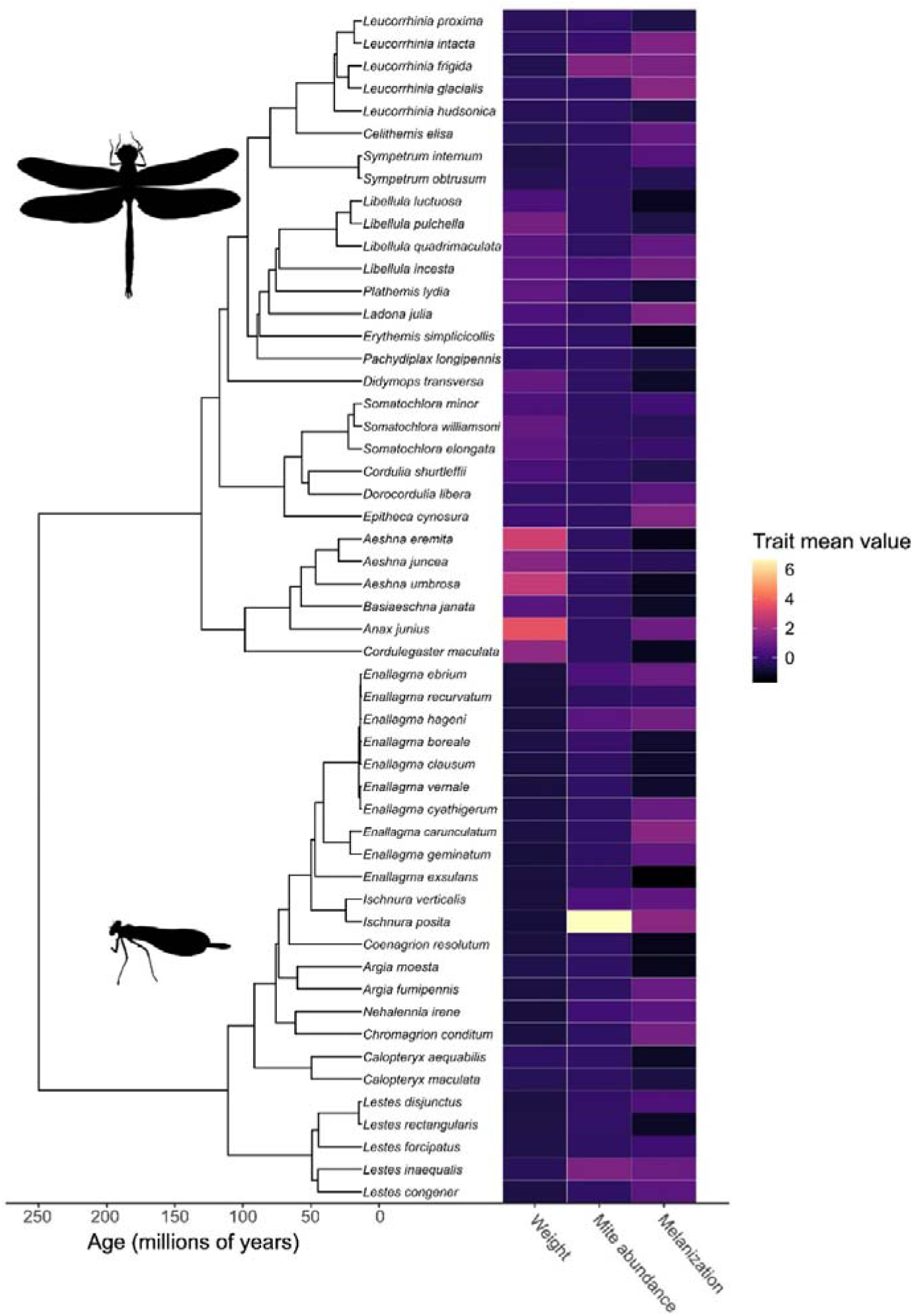
Phylogenetic tree of the species used in our analyses, with trait data on body weight, mite abundance, and melanization mapped onto the tree. All three traits are mean-centered and scaled, such that they have a mean of 0 and SD of 1, with darker colors denoting lower values and lighter colors denoting higher values. Inset images of the dragonfly and damselfly represent the two suborders within the clade.

**Fig 5.**
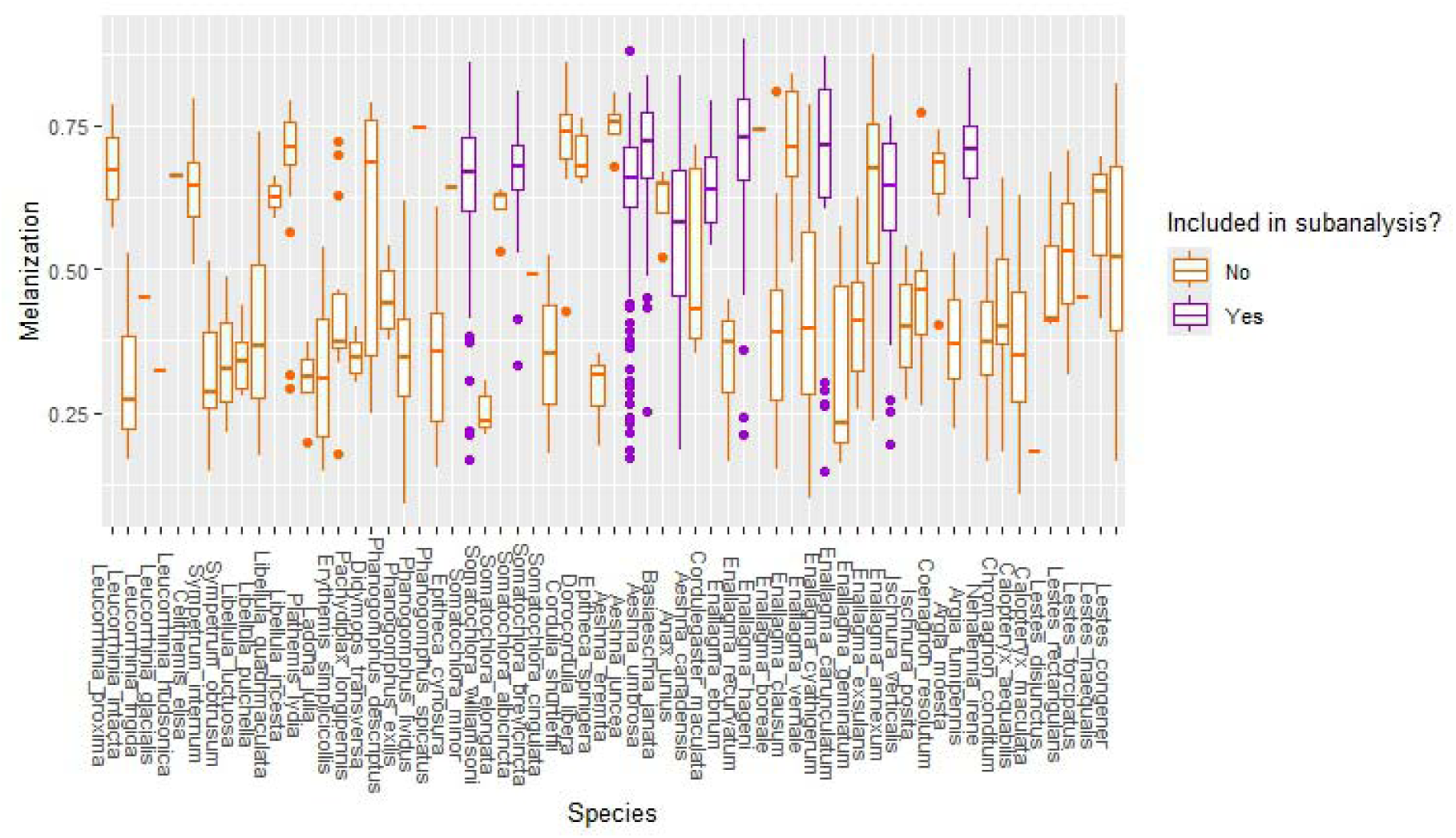
Boxplots representing the variation in melanization response (y-axis) of the various species (x-axis) included in our analyses. Color denotes those species included in the sub-analysis using SEMs.

## Discussion

Immune responses are necessary to defend organisms from the deleterious and fitness-reducing effects of parasites (and disease in general, Hasik & Siepielski 2022a). Although there is increasing interest in the macroecology of parasites and infectious diseases (Stephens *et al*. 2016), we know very little about the macroecology of immune responses (LoScerbo *et al*. 2020). Understanding geographic and among-population variation in immune defenses is key to predicting the impact of parasites on host populations, but variation in immune defenses themselves rely on myriad factors (Becker *et al*. 2019). In this study we used data from a widely distributed and well-studied host-parasite system to test the hypothesis that immune defenses respond to geographical variation in abiotic and biotic factors. Contrary to our expectations, the most prominent result is that the abiotic environment, rather than parasitism or host characteristics, best predicted geographic variation in melanization response. Structural equation models applied to a subset of co-distributed species further supported these findings, with additional insight into the influence of parasitism. Overall, our results are consistent with the strength of melanization being a by-product of adaptation to desiccation and to some degree an evolutionary response to the abundance of parasites in the environment.

Our results add to a growing body of literature showing how environmental conditions set the stage for parasitism, in this case via the relationship between environmental factors and immune defenses. Evidence from mammals (Hasik *et al*. 2024b), insects (LoScerbo *et al*. 2020; Hasik & Siepielski 2022b; Hasik *et al*. 2024a), fish (Bolnick *et al*. 2020), and plants (Penczykowski *et al*. 2014) has revealed that abiotic and biotic factors predict parasitism due to the varied connections between these factors and their effects on the host and/or parasites. In the case of odonates and their water mite parasites, water pH is a key driver of variation in odonate host community composition (Arrowsmith *et al*. 2018) and water mite infection levels (LoScerbo *et al*. 2020; Hasik & Siepielski 2022b; Hasik *et al*. 2024a). The suggested mechanism by which water pH (acidic conditions) affects parasitism is a reduction in water mite larval survival and egg viability at low pH (Edwards 2004). However, the results from our SEM revealed that, at least in the populations and species that we sampled, pH is neither a predictor of melanization nor mite parasitism. Instead, precipitation and to some degree temperature were the primary drivers of melanization. The reasons for this are unclear, and mean annual temperature may not necessarily be a good indicator of what the temperature was at the time of sampling, but it could relate to how the local environment is expected to affect melanization in insects.

Climatic conditions best explained variation in the strength of immune response among host individuals and populations. One possible reason behind this pattern is that abiotic factors directly contribute to immunity in ectotherms (Ferguson *et al*. 2018). In general, high temperatures drive greater melanization responses (Fedorka *et al*. 2013). Melanin production is primarily used for three functions: parasite defense (as is the case against mites, Hasik *et al*. 2023), thermoregulation via cuticle darkening (Nakhleh *et al*. 2017), and water retention via reduction in cuticular permeability (King & Sinclair 2015). These functions may covary inversely due to a resource allocation trade-off and depending on how much they are being selected for (Cotter *et al*. 2008).

Our results emphasize the strong influence of annual precipitation on melanization response, with the greatest responses recorded in the driest regions. To our knowledge, this is the first study to document a relationship between immune response and precipitation and/or humidity. Nevertheless, some studies have shown that cuticle darkening driven by high cuticular melanin content is affected by how humid an environment is, yet the evidence is equivocal (Clusella-Trullas & Nielsen 2020; Klunk *et al*. 2022). Similarly, the degree of cuticle darkening varies geographically among wild populations of *D. melanogaster* from Africa (Pool & Aquadro 2006), India (Parkash *et al*. 2008), and Australia (Telonis-Scott *et al*. 2011), which indicate that cuticle darkness is an adaptation to regional climatic conditions. Laboratory experiments with *D. melanogaster* further indicate that individuals with darker cuticles are more resistant to desiccation (Ramniwas *et al*. 2013), which supports the melanization-desiccation hypothesis. The observed increase in desiccation resistance is likely due to a decrease in cuticular permeability. Taken together, these results suggest that the strong melanin encapsulation response observed in the driest part of our study region could be a by-product of the adaptive value of melanin for resistance to desiccation. These results contrast with the common expectation that because wet habitats contain more parasites and predators, host cuticles should exhibit higher cuticle melanization (Delhey 2019; Clusella-Trullas & Nielsen 2020). In such a case, wet environments could select for cuticles that are more heavily melanized at the expense of the immune defenses.

Across all individuals, we found that there was no overall relationship between degree of parasitism and melanization response. The reasons for this in our overall analysis are unclear; however, our interpopulation comparisons using SEMs and a subset of our overall data suggested that local adaptation of the hosts to their water mite parasites may be occurring. That is, the melanization-based immune defense could increase with the degree of local adaptation of odonate hosts to their parasites (Gómez-Llano *et al*. 2020). Indeed, hosts from populations with greater parasite abundance had a stronger melanization response, independent of the negative relationship we found between individual deviance and melanization, supporting previous work showing that the melanin-based immune defense used in this study is greater in populations experiencing heavy parasitism (Nagel *et al*. 2010; Kaunisto *et al*. 2017). This suggests that hosts from high-risk populations have greater melanization response, which could be an adaptive response, but heavy infections could be depleting or suppressing melanin stores and thereby reducing the melanization response. The latter hypothesis is highly parsimonious given the mechanics of the melanization response. That is, odonates produce melanin to seal mite feeding tubes and defend themselves from attack (Hasik *et al*. 2023). Heavy infections would stimulate the phenoloxidase cascade to produce more melanin than would be seen in a lighter infection, hence the negative relationship with individual deviance.

Our analysis testing for a phylogenetic signal of various traits of interest revealed very few significant patterns. Specifically, we only found a phylogenetic signal in body weight, which is consistent with previous work (LoScerbo *et al*. 2020). The lack of a signal for both mite abundance and immune defenses bolster our other findings that both of these traits of interest are mediated by the environment and are not constrained by phylogeny, suggesting that accounting for species effects are more important than accounting for phylogeny in analyses of odonate immune defenses. While we did not study evolutionary mechanisms *per se*, such species-specific variation highlights how environmental differences among host-parasite populations may result in evolutionary hot- and cold-spots (Thompson 2005). That is, our finding that environmental variation best explains variation in immunity, previous findings relating the local environment to water mite parasitism (LoScerbo *et al*. 2020; Hasik & Siepielski 2022b; Hasik *et al*. 2024a), and the lack of signal in our phylogenetic analyses, suggest that neither the host nor parasite are the primary drivers of the (co)evolutionary relationships. Instead, the local environment shapes these interactions, after which the host and parasite respond. Indeed, the Stockholm Paradigm posits that ecological disruptions generate novel combinations of host and parasite assemblages (Brooks *et al*. 2015), and it is these novel associations that then drive host-parasite (co)evolution.

Our study is based on a wealth of field data, including data gained from over one thousand immune assays, but we must mention a number of important caveats. First, all our analyses are based on correlational data. Though our modeling framework utilizing SEMs allows for tests of hypothesized “casual” pathways, they cannot infer causality. Second, we used a general measure of parasitism and did not identify water mites to the species level. *Arrenurus* is a remarkably diverse subgenus (Więcek *et al*. 2023), and while we did not find evidence for parasites being a primary driver of immune defenses there remains the possibility that such relationships exist on a more species-specific basis. Finally, though our overall dataset contained data collected from 63 species of odonate hosts, some species had better representation than others. Collecting more data from the larger, less-represented species such as *Aeshna* dragonflies is necessary to understand if the patterns revealed in our study are representative of what was found for the smaller, well-represented species such as *Ischnura* and *Enallagma* damselflies.

In conclusion, our results emphasize the role of the local environment in mediating host-parasite interactions. Because the factors driving variation in immunity among individuals and populations are poorly-studied and rarely factored into epidemiologic models (Becker *et al*. 2019), the findings of such models may be biased. Our study demonstrates that analyzing how the climate and environmental factors affect immune defenses reveals nuance inherent to host-parasite interactions. Host characteristics, parasites, and the abiotic environment are known to affect immunity in complex ways (Sweeny & Albery 2022). Understanding and disentangling these relationships can be difficult (Johnson *et al*. 2019), but future studies using approaches similar to those employed here can improve our knowledge of the emergent differences in parasitism among host individuals and species. Finally, our findings may also be used to understand host-parasite relationships under climate change scenarios, specifically how temperature and precipitation may contribute to patterns of immune defense. Given the synergistic effects of these two factors, increases in temperature or reduction in precipitation at northern latitudes will also influence parasitism rate. Future experimental studies should clarify the nature of this effect.

## Acknowledgements

Concordia University is located on unceded Indigenous lands. The Kanien’kehá:ka Nation is recognized as the custodians of Tiohtià:ke/Montreal. We thank Serena Sinno, Gabriel Muñoz, and Javier Ibarra-Isassi for help and advice throughout the project development. AZH benefitted from the musical inspiration of Spiritbox. This research project was funded by a NSERC Discovery Grant (RGPINLJ2015LJ06081) to JPL.

## Author contributions

JPL conceived the study. AZH, MB, EJP, and JPL designed the study and analyses. MB, JH, TG, and JPL collected field data, MB, JH, TG and TdB processed samples in the lab. AZH performed modelling work and analyzed data. AZH wrote the first draft of the manuscript, and all authors contributed substantially to revisions. JPL acquired funding. Our study brings together authors from multiple countries, including scientists based in the country where the study was carried out.

