## Supplementary Information for "The macroecology of immunity: predominant influence of climate on invertebrate immune response"

**Figures S1-S3, Table S1, details on the immune assays**

**Fig. S1** –
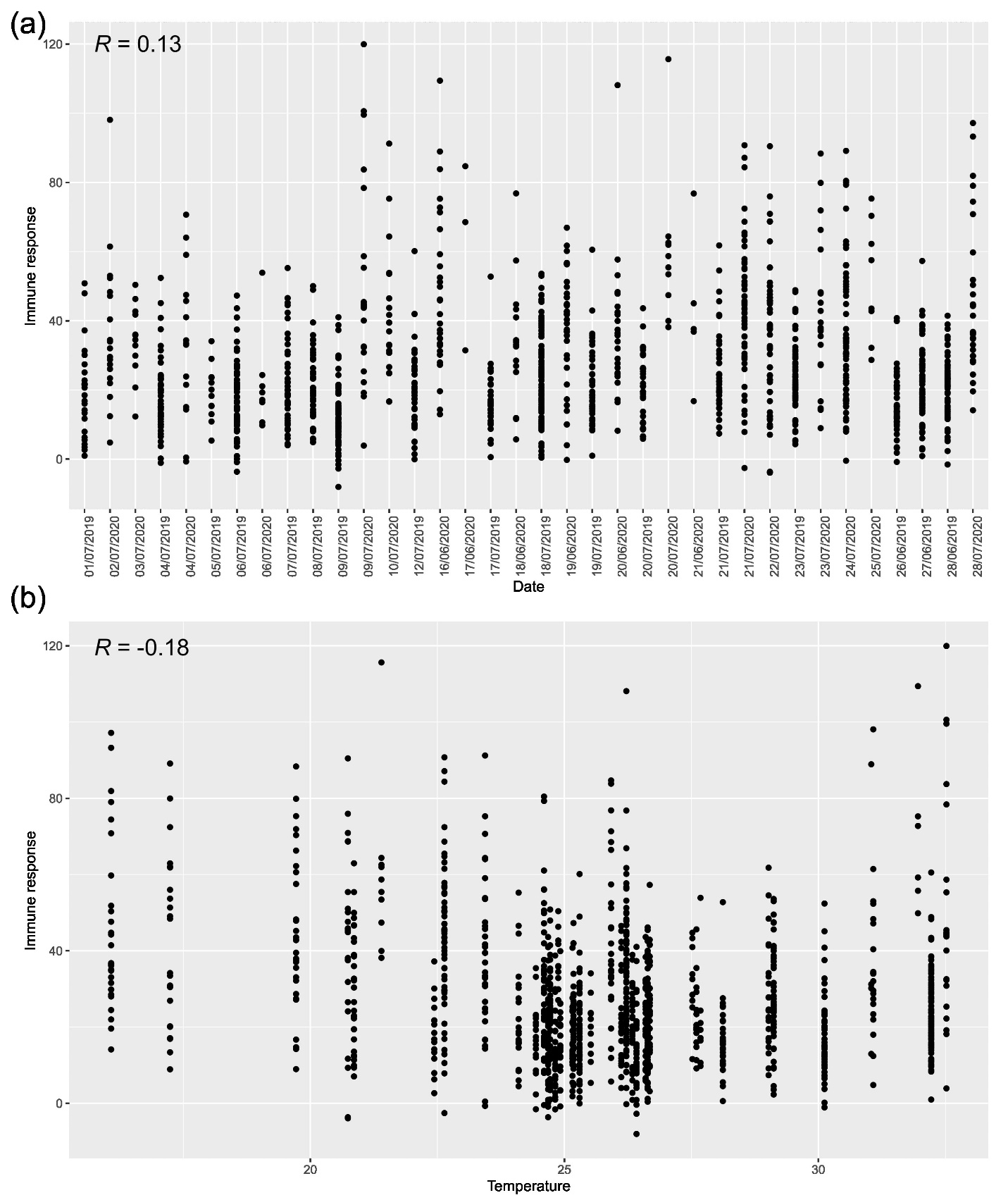
 No evidence for an effect of temperature at time of collection or date (i.e., seasonal effects) on immune responses. The x-axes denote date of sampling (a) and temperature at time of collection (b), with immune response on the y-axis of both plots. Inset text denote the correlation coefficients.

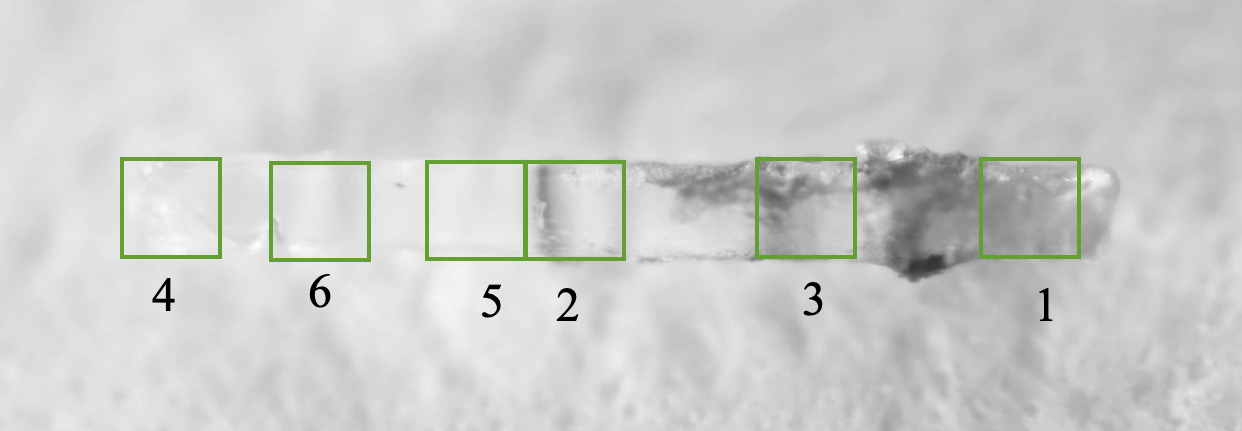
**Fig. S2** - Illustration of regions that were measured to quantify darkness values.

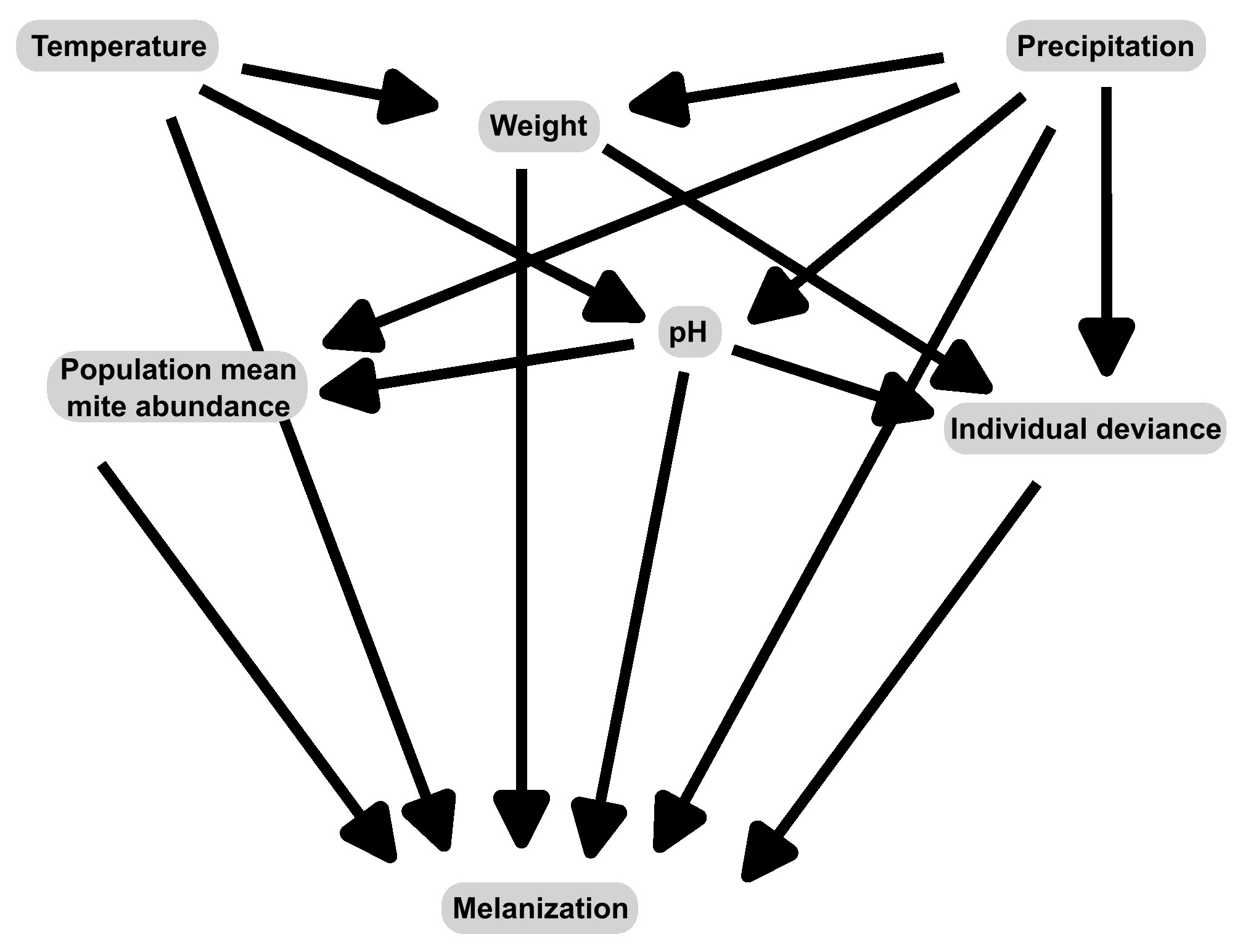
**Fig. S3** – Graphical representation of the initial modeling framework used in our structural equation models. Directed arrows denote the influence of a predictor on a response variable.

**Table S1 -** Summary information on the species and samples sizes collected at each site.

| Site | Species | *n* |
| --- | --- | --- |
| 1000 Iles 1 | *Argia moesta* | 1 |
|  | *Enallagma annexum* | 3 |
|  | *Ischnura verticalis* | 1 |
|  | *Lestes disjunctus* | 9 |
|  | *Lestes rectangularis* | 6 |
|  | *Sympetrum obtrusum* | 14 |
| Bouzan | *Cordulia shurtleffii* | 1 |
|  | *Dorocordulia libera* | 4 |
|  | *Enallagma ebrium* | 4 |
|  | *Leucorrhinia frigida* | 1 |
|  | *Leucorrhinia proxima* | 1 |
|  | *Nehalennia irene* | 1 |
| Brousseau | *Chromagrion conditum* | 8 |
|  | *Enallagma ebrium* | 7 |
|  | *Enallagma hageni* | 9 |
|  | *Epitheca cynosura* | 3 |
|  | *Ischnura posita* | 4 |
|  | *Ischnura verticalis* | 7 |
|  | *Ladona julia* | 9 |
|  | *Leucorrhinia intacta* | 2 |
|  | *Nehalennia irene* | 5 |
| Brown | *Calopteryx maculata* | 7 |
|  | *Enallagma ebrium* | 1 |
|  | *Enallagma hageni* | 11 |
|  | *Ischnura verticalis* | 4 |
|  | *Ladona julia* | 1 |
|  | *Lestes inaequalis* | 2 |
|  | *Leucorrhinia frigida* | 26 |
|  | *Leucorrhinia proxima* | 1 |
|  | *Libellula incesta* | 9 |
| Bruno 2 | *Anax junius* | 1 |
|  | *Enallagma annexum* | 1 |
|  | *Enallagma ebrium* | 1 |
|  | *Erythemis simplicicollis* | 1 |
|  | *Libellula incesta* | 2 |
|  | *Libellula luctuosa* | 3 |
|  | *Libellula pulchella* | 6 |
|  | *Pachydiplax longipennis* | 4 |
|  | *Plathemis lydia* | 1 |
|  | *Sympetrum internum* | 1 |
| Bruyere | *Enallagma hageni* | 18 |
|  | *Epitheca spinigera* | 13 |
|  | *Libellula quadrimaculata* | 2 |
|  | *Phanogomphus spicatus* | 7 |
| Drolet | *Cordulia shurtleffii* | 1 |
|  | *Dorocordulia libera* | 1 |
|  | *Enallagma ebrium* | 11 |
|  | *Ladona julia* | 3 |
|  | *Leucorrhinia frigida* | 4 |
|  | *Phanogomphus spicatus* | 2 |
| Etang John | *Enallagma clausum* | 2 |
|  | *Enallagma vernale* | 1 |
|  | *Ischnura verticalis* | 3 |
| Fen Delage | *Aeshna umbrosa* | 1 |
|  | *Chromagrion conditum* | 1 |
|  | *Dorocordulia libera* | 1 |
|  | *Ladona julia* | 3 |
|  | *Leucorrhinia frigida* | 8 |
|  | *Leucorrhinia hudsonica* | 2 |
|  | *Leucorrhinia proxima* | 2 |
|  | *Nehalennia irene* | 3 |
| Gale | *Chromagrion conditum* | 4 |
|  | *Ischnura verticalis* | 8 |
|  | *Ladona julia* | 3 |
|  | *Phanogomphus exilis* | 12 |
| Gamble | *Enallagma ebrium* | 10 |
|  | *Enallagma hageni* | 6 |
|  | *Libellula quadrimaculata* | 1 |
| Gauthier | *Calopteryx maculata* | 2 |
|  | *Enallagma hageni* | 2 |
|  | *Ischnura verticalis* | 3 |
|  | *Lestes disjunctus* | 6 |
|  | *Lestes forcipatus* | 4 |
|  | *Leucorrhinia proxima* | 3 |
|  | *Somatochlora elongata* | 2 |
|  | *Sympetrum obtrusum* | 6 |
| Hidden | *Argia fumipennis* | 2 |
|  | *Celithemis elisa* | 2 |
|  | *Enallagma ebrium* | 5 |
|  | *Enallagma hageni* | 3 |
|  | *Ischnura verticalis* | 2 |
|  | *Libellula incesta* | 1 |
| Ile d l Vis | *Argia moesta* | 6 |
|  | *Enallagma exsulans* | 6 |
| LDS | *Enallagma carunculatum* | 1 |
|  | *Enallagma ebrium* | 15 |
|  | *Ladona julia* | 1 |
|  | *Leucorrhinia proxima* | 1 |
|  | *Libellula quadrimaculata* | 2 |
| Libby | *Calopteryx maculata* | 1 |
|  | *Enallagma hageni* | 8 |
|  | *Ischnura verticalis* | 6 |
|  | *Ladona julia* | 3 |
| Long Pond | *Enallagma ebrium* | 2 |
|  | *Ischnura verticalis* | 13 |
|  | *Nehalennia irene* | 1 |
|  | *Phanogomphus exilis* | 3 |
|  | *Phanogomphus spicatus* | 1 |
| Marais N | *Enallagma ebrium* | 4 |
|  | *Ladona julia* | 2 |
|  | *Lestes congener* | 1 |
|  | *Lestes disjunctus* | 4 |
|  | *Leucorrhinia hudsonica* | 1 |
|  | *Leucorrhinia proxima* | 2 |
|  | *Libellula quadrimaculata* | 1 |
|  | *Nehalennia irene* | 3 |
|  | *Somatochlora williamsoni* | 1 |
|  | *Sympetrum obtrusum* | 11 |
| Marise | *Aeshna canadensis* | 1 |
|  | *Enallagma hageni* | 10 |
|  | *Lestes disjunctus* | 1 |
|  | *Lestes inaequalis* | 1 |
|  | *Leucorrhinia frigida* | 12 |
|  | *Leucorrhinia glacialis* | 1 |
|  | *Libellula incesta* | 1 |
|  | *Nehalennia irene* | 1 |
| Marlon | *Chromagrion conditum* | 10 |
|  | *Dorocordulia libera* | 7 |
|  | *Enallagma ebrium* | 2 |
|  | *Leucorrhinia hudsonica* | 1 |
|  | *Nehalennia irene* | 5 |
|  | *Phanogomphus spicatus* | 11 |
| MDS | *Aeshna eremita* | 1 |
|  | *Coenagrion resolutum* | 5 |
|  | *Enallagma boreale* | 8 |
|  | *Enallagma ebrium* | 1 |
|  | *Leucorrhinia hudsonica* | 2 |
|  | *Leucorrhinia proxima* | 1 |
|  | *Libellula quadrimaculata* | 6 |
|  | *Somatochlora albicincta* | 5 |
| Mercier 1 | *Cordulegaster maculata* | 12 |
|  | *Cordulia shurtleffii* | 3 |
|  | *Enallagma boreale* | 8 |
|  | *Leucorrhinia hudsonica* | 7 |
|  | *Leucorrhinia proxima* | 7 |
|  | *Libellula quadrimaculata* | 4 |
|  | *Somatochlora brevicincta* | 11 |
|  | *Somatochlora cingulata* | 1 |
|  | *Somatochlora elongata* | 1 |
| Moore | *Cordulia shurtleffii* | 1 |
|  | *Dorocordulia libera* | 3 |
|  | *Enallagma ebrium* | 2 |
|  | *Enallagma hageni* | 1 |
|  | *Ladona julia* | 27 |
|  | *Phanogomphus spicatus* | 5 |
| Muskrat | *Argia fumipennis* | 22 |
|  | *Enallagma geminatum* | 4 |
|  | *Ischnura verticalis* | 4 |
|  | *Lestes inaequalis* | 2 |
| Noranda | *Enallagma boreale* | 1 |
|  | *Enallagma ebrium* | 9 |
|  | *Enallagma hageni* | 6 |
|  | *Epitheca spinigera* | 1 |
|  | *Ladona julia* | 6 |
|  | *Leucorrhinia intacta* | 1 |
|  | *Libellula quadrimaculata* | 4 |
|  | *Phanogomphus spicatus* | 16 |
| Orford | *Enallagma boreale* | 1 |
|  | *Enallagma clausum* | 1 |
|  | *Enallagma cyathigerum* | 1 |
|  | *Enallagma ebrium* | 4 |
|  | *Enallagma hageni* | 1 |
|  | *Ischnura verticalis* | 7 |
|  | *Leucorrhinia intacta* | 1 |
| Orignal | *Enallagma ebrium* | 2 |
|  | *Enallagma hageni* | 10 |
|  | *Leucorrhinia frigida* | 8 |
|  | *Libellula incesta* | 4 |
| Osisko | *Enallagma ebrium* | 1 |
|  | *Ischnura verticalis* | 27 |
| Parker | *Chromagrion conditum* | 3 |
|  | *Enallagma ebrium* | 2 |
|  | *Enallagma hageni* | 1 |
|  | *Ischnura verticalis* | 12 |
|  | *Ladona julia* | 9 |
|  | *Leucorrhinia intacta* | 1 |
|  | *Nehalennia irene* | 2 |
|  | *Phanogomphus exilis* | 2 |
| Pelletier | *Enallagma ebrium* | 3 |
|  | *Enallagma hageni* | 5 |
|  | *Ischnura verticalis* | 20 |
|  | *Leucorrhinia intacta* | 1 |
|  | *Libellula quadrimaculata* | 22 |
| Pike 2 | *Argia moesta* | 13 |
|  | *Basiaeschna janata* | 3 |
|  | *Calopteryx aequabilis* | 3 |
|  | *Calopteryx maculata* | 23 |
|  | *Didymops transversa* | 4 |
|  | *Phanogomphus descriptus* | 9 |
|  | *Phanogomphus exilis* | 1 |
|  | *Phanogomphus lividus* | 7 |
| Puant | *Aeshna canadensis* | 1 |
|  | *Enallagma ebrium* | 13 |
|  | *Enallagma hageni* | 2 |
|  | *Lestes congener* | 4 |
|  | *Lestes disjunctus* | 37 |
|  | *Lestes forcipatus* | 1 |
|  | *Leucorrhinia frigida* | 1 |
|  | *Leucorrhinia proxima* | 2 |
|  | *Nehalennia irene* | 5 |
|  | *Sympetrum internum* | 2 |
|  | *Sympetrum obtrusum* | 16 |
| R Epaule | *Aeshna eremita* | 6 |
|  | *Coenagrion resolutum* | 4 |
|  | *Cordulia shurtleffii* | 8 |
|  | *Enallagma boreale* | 6 |
|  | *Enallagma ebrium* | 1 |
|  | *Ischnura verticalis* | 1 |
|  | *Lestes disjunctus* | 4 |
|  | *Leucorrhinia hudsonica* | 2 |
|  | *Leucorrhinia proxima* | 7 |
|  | *Libellula quadrimaculata* | 3 |
| R Montmo | *Aeshna juncea* | 1 |
|  | *Calopteryx aequabilis* | 5 |
|  | *Cordulegaster maculata* | 7 |
|  | *Enallagma boreale* | 1 |
|  | *Leucorrhinia hudsonica* | 1 |
|  | *Somatochlora albicincta* | 3 |
|  | *Somatochlora minor* | 4 |
| Rapides | *Enallagma exsulans* | 4 |
|  | *Ischnura verticalis* | 4 |
| Riv Sud 1 | *Epitheca cynosura* | 1 |
|  | *Ischnura verticalis* | 3 |
| Riv Sud 2 | *Ischnura verticalis* | 6 |
|  | *Plathemis lydia* | 3 |
| SDLN 1 | *Chromagrion conditum* | 1 |
|  | *Cordulia shurtleffii* | 1 |
|  | *Dorocordulia libera* | 6 |
|  | *Enallagma clausum* | 2 |
|  | *Enallagma vernale* | 1 |
|  | *Ischnura verticalis* | 1 |
|  | *Leucorrhinia proxima* | 4 |
|  | *Libellula luctuosa* | 1 |
|  | *Libellula pulchella* | 3 |
|  | *Libellula quadrimaculata* | 5 |
|  | *Plathemis lydia* | 3 |
| St Bruno | *Enallagma ebrium* | 2 |
|  | *Erythemis simplicicollis* | 6 |
|  | *Ischnura verticalis* | 1 |
|  | *Leucorrhinia intacta* | 1 |
|  | *Libellula incesta* | 3 |
|  | *Libellula luctuosa* | 5 |
|  | *Libellula pulchella* | 2 |
| St Charles | *Calopteryx maculata* | 2 |
|  | *Enallagma ebrium* | 4 |
|  | *Enallagma recurvatum* | 1 |
|  | *Ladona julia* | 1 |
|  | *Plathemis lydia* | 1 |
|  | *Sympetrum obtrusum* | 1 |
| Surprise | *Aeshna canadensis* | 1 |
|  | *Dorocordulia libera* | 1 |
|  | *Enallagma hageni* | 3 |
|  | *Lestes disjunctus* | 1 |
|  | *Lestes inaequalis* | 1 |
|  | *Leucorrhinia frigida* | 21 |
|  | *Libellula incesta* | 12 |
|  | *Nehalennia irene* | 1 |
| Waterloo | *Enallagma ebrium* | 19 |
|  | *Enallagma hageni* | 1 |
|  | *Epitheca cynosura* | 5 |
|  | *Ischnura verticalis* | 9 |

**Details on the immune assays**

We placed adult odonates in a manila envelope after capture and brought them back to the lab. Individuals spent no more than 2 hours in an envelope from the time they were caught to the moment they were placed in a mesh box to acclimate. The mesh boxes were ~ 11cm × 11cm × 11cm, and included a wet paper towel at the bottom.  We gave the odonates 30 minutes to acclimate (Combes *et al.* 2012) at room temperature with full natural lighting (and all lights turned on in the lab) before beginning the experiment, and conducted all experiments at room temperature (~20-23 °C) (Murdock *et al.* 2012).

To perform the immune response assay, we inserted a sterilized 3mm piece of nylon monofilament (hereafter implant) into the venter of the odonate thorax. This region of the body was chosen because it is where mites tend to latch onto their host, and the implant is meant to imitate the feeding tube of the parasite (Hasik *et al.* 2023). After insertion, we returned odonates to their breathable container with a wet paper towel for 12 hours to allow melanization of the implant (after ~8 hours no more melanin is allocated to an implant in an odonate adult; Contreras-Garduño *et al.* 2006). We did not provide a food source during these 12 hours, and subjected odonates to their typical photoperiod via natural lighting. After 12 hours we removed and photographed the implant directly above (~30cm between implant and camera lens, centering the lens over the implant) using the flash from the camera (Canon EOS Rebel T6s, lens extension MP-E 65mm). We then photographed the complete odonate animal, sacrificed them with acetone, and then stored them in glassine envelopes for subsequent identification and mite counting.

**Details on immune quantification**

We used six measurement points because it was necessary to ensure that the areas measured along the filament were standardized. This differs from the method used in Ilvonen & Suhonen (2016), where the upper and lower portions on both the sides of the nylon implant were measured (giving four measurements total). However, this method could not work for our study since the length of the portion inserted and the portion not inserted are not constant between and within images, as species varied markedly in size. Placing an insert into a damselfly to the same length that we did into a dragonfly would result in it penetrating the dorsal side of the thorax. By using three square areas on either side of the insertion point of the nylon we were able to keep the area measured on each side of the insertion point consistent within a given image, and proportional between images. In addition, we chose six areas because the melanization was not constant along the entirety of the inserted region of the filament, and taking three measurements along this region allowed us to measure the extreme ends and middle of ends of the insert. Finally, we compared the average grey value of the three measurements spanning the inserted portion to the average grey value of the three measurements spanning the portion that was not inserted were taken.

1.

Combes, S.A., Rundle, D.E., Iwasaki, J.M. & Crall, J.D. (2012). Linking biomechanics and ecology through predator–prey interactions: flight performance of dragonflies and their prey. *Journal of Experimental Biology*, 215, 903-913.

2.

Contreras-Garduño, J., Canales-Lazcano, J. & Córdoba-Aguilar , A. (2006). Wing pigmentation, immune ability, fat reserves and territorial status in males of the rubyspot damselfly, *Hetaerina americana*. *Journal of Ethology*, 24, 165-173.

3.

Hasik, A.Z., Ilvonen, J.J., Siepielski, A.M. & Murray, R.L. (2023). Odonata immunity, pathogens, and parasites. In: *Dragonflies and Damselflies* (eds. Córdoba-Aguilar, A, Beatty, CD & Bried, JT). Oxford University Press.

4.

Ilvonen, J.J. & Suhonen, J. (2016). Phylogeny affects host's weight, immune response and parasitism in damselflies and dragonflies. 33, 160421.

5.

Murdock, C.C., Paaijmans, K.P., Bell, A.S., King, J.G., Hillyer, J.F., Read, A.F. *et al.* (2012). Complex effects of temperature on mosquito immune function. *Proceedings of the Royal Society B*, 279, 20120638.
